# EBNA1 SUMOylation by PIAS1 Suppresses EBV Lytic Replication and Enhances Episome Maintenance

**DOI:** 10.1101/2025.09.01.673444

**Authors:** Febri Gunawan Sugiokto, Kun Zhang, Renfeng Li

## Abstract

Epstein-Barr virus nuclear antigen 1 (EBNA1) is essential for the replication and stable maintenance of the viral episome in infected cells. Here, we identify the SUMO E3 ligase PIAS1 as a key regulator of EBNA1 through site-specific SUMOylation. Our Chromatin-Immunoprecipitation Sequencing (ChIP-seq) analysis revealed that PIAS1 is enriched at the viral origin of plasmid replication (*oriP*), where it physically associates with EBNA1 and catalyzes its SUMOylation. Using mutational analysis, we identified three lysine residues on EBNA1 (K17, K75, and K241) as major SUMOylation sites. Disruption of these sites compromises EBNA1’s ability to restrict EBV lytic replication. In addition, both PIAS1 depletion and the disruption of EBNA1 SUMOylation lead to reduced retention of EBNA1-*OriP*-based EBV mini-replicon, indicating the importance of EBNA1 SUMOylation in viral episome maintenance. Together, these results uncover a conserved post-translational mechanism by which PIAS1-mediated SUMOylation modulates EBNA1 function and EBV episome maintenance and suggests a broader role for SUMOylation in viral latency, lytic replication and persistence.

**IMPORTANCE:** Epstein-Barr virus (EBV) persists in infected cells by maintaining its episome through the viral protein EBNA1. We discovered that PIAS1 SUMOylates EBNA1 at specific sites, a process essential for EBNA1 to retain viral episome and suppress reactivation. When SUMOylation is disrupted, EBV-based replicon becomes less stable and EBV is more likely to reactivate. These findings reveal a new layer of host control of EBV latency and reactivation, and highlight PIAS1-mediated EBNA1 SUMOylation as a key mechanism regulating viral persistence.

## INTRODUCTION

Epstein-Barr virus (EBV), a ubiquitous human gamma-herpesvirus, establishes lifelong persistent infection primarily in memory B cells. While often asymptomatic, EBV infection is linked to a range of malignancies, including endemic Burkitt’s lymphoma, nasopharyngeal carcinoma, Hodgkin lymphoma, and a subset of gastric carcinomas. During latency, EBV expresses a restricted set of viral genes that facilitate viral genome persistence, immune evasion, and host cell survival [1–3].

As a central player in EBV latency, EBNA1 is essential for the replication and maintenance of the viral episome during cell division. EBNA1 binds to the dyad symmetry (DS) and family of repeats (FR) elements within the viral origin of plasmid replication (*oriP*), coordinating both the replication and mitotic segregation of the EBV genome. In addition to its role in episome maintenance, EBNA1 regulates the transcription of both viral and cellular genes. Structurally, EBNA1 contains a C-terminal DNA-binding and dimerization domain required for *oriP* interaction, and an N-terminal glycine-alanine repeat (GAr) region that limits proteasomal degradation and antigen presentation, thereby contributing to immune evasion [3–8].

In addition to EBV, other DNA viruses such as Kaposi’s sarcoma-associated herpesvirus (KSHV) and human papillomavirus (HPV) also maintain their genomes as episomes during latent or persistent infection [9]. These viruses also encode episome maintenance proteins, namely LANA in KSHV and E2 in HPV, that perform analogous functions critical for viral genome persistence. LANA tethers the KSHV episome to host chromatin through interactions with histones and chromosomal proteins such as BRD2/4, while also recruiting cellular replication machinery to ensure genome maintenance [10–13]. Similarly, the E2 protein of HPV binds to specific sequences in the viral origin and interacts with host mitotic chromosomes, via Brd4 and TopBP1, to facilitate episome segregation during cell division [14, 15]. Like EBNA1, both LANA and E2 are also involved in regulating viral transcription and replication. This functional conservation highlights a shared evolutionary strategy among diverse viruses to ensure long-term persistence in host cells through episome tethering, DNA replication, and transcriptional control [9].

SUMOylation is a highly regulated post-translational modification (PTM) process involving the covalent attachment of Small Ubiquitin-like Modifier (SUMO) proteins— SUMO1, SUMO2, and SUMO3 in humans—to lysine residues on target proteins. This reversible modification is crucial for various cellular processes, including transcriptional regulation, DNA repair, nuclear-cytosolic transport, signal transduction, and protein stability. The SUMOylation pathway comprises a well-coordinated enzymatic cascade, initiated by the activation of SUMO precursors through the E1 activating enzyme complex (SAE1 and SAE2, also known as UBA2). Subsequently, SUMO is transferred to the E2 conjugating enzyme UBC9, which uniquely serves as the sole E2 enzyme responsible for SUMO transfer in mammals [16].

While UBC9 can mediate SUMO conjugation independently, E3 ligases significantly enhance substrate specificity and conjugation efficiency. The Protein Inhibitor of Activated STAT (PIAS) family proteins, comprising PIAS1, PIAS2 (also known as PIASx), PIAS3, and PIAS4 (also known as PIASy), play an important role among these E3 ligases. PIAS proteins facilitate the transfer of SUMO from UBC9 to substrate proteins by target selection, which could influence the function and the subcellular localization of SUMOylated targets. These ligases are subject to regulation and have been implicated in various biological processes, such as immune response, genome stability, and the control of oncogenic signaling pathways [17].

Our previous research has established PIAS1 as an EBV restriction factor and an E3 ligase that synergizes with SAMHD1 and YTHDF2 to regulate EBV replication through SUMOylation [18–22]. In this study, we demonstrated that PIAS1 is localized in EBV *oriP*, where it interacts with EBNA1. We found that PIAS1 enhances EBNA1 SUMOylation at three specific lysine residues: K17, K75, and K241. The interaction between EBNA1 and PIAS1, along with the PIAS1-mediated SUMOylation process, plays an important role in regulating viral replication and genome maintenance.

## RESULTS

### PIAS1 is enriched at EBV *oriP* in EBV+ lymphoma cells

In our previous study, we demonstrated that PIAS1 plays a crucial role in restricting EBV lytic replication. Specifically, PIAS1 inhibits lytic viral gene transcription by binding to viral promoters [20]. To gain further insight into PIAS1 binding across the entire EBV genome, we conducted ChIP-Seq experiment using Akata (EBV+) Burkitt lymphoma cells as our model system. Unexpectedly, we observed that PIAS1 peaks were significantly enriched at the EBV *oriP* (FR and DS) region (**Fig. 1A and B**).

To validate these findings, we performed ChIP-qPCR, which confirmed the enrichment of PIAS1 at the EBV *oriP* with anti-PIAS1 antibody compared to the IgG control (**Fig. 1C**). These results suggest PIAS1 may regulate EBV replication and genome maintenance, potentially through interaction with the viral *oriP* or EBNA1.

**Figure 1.**
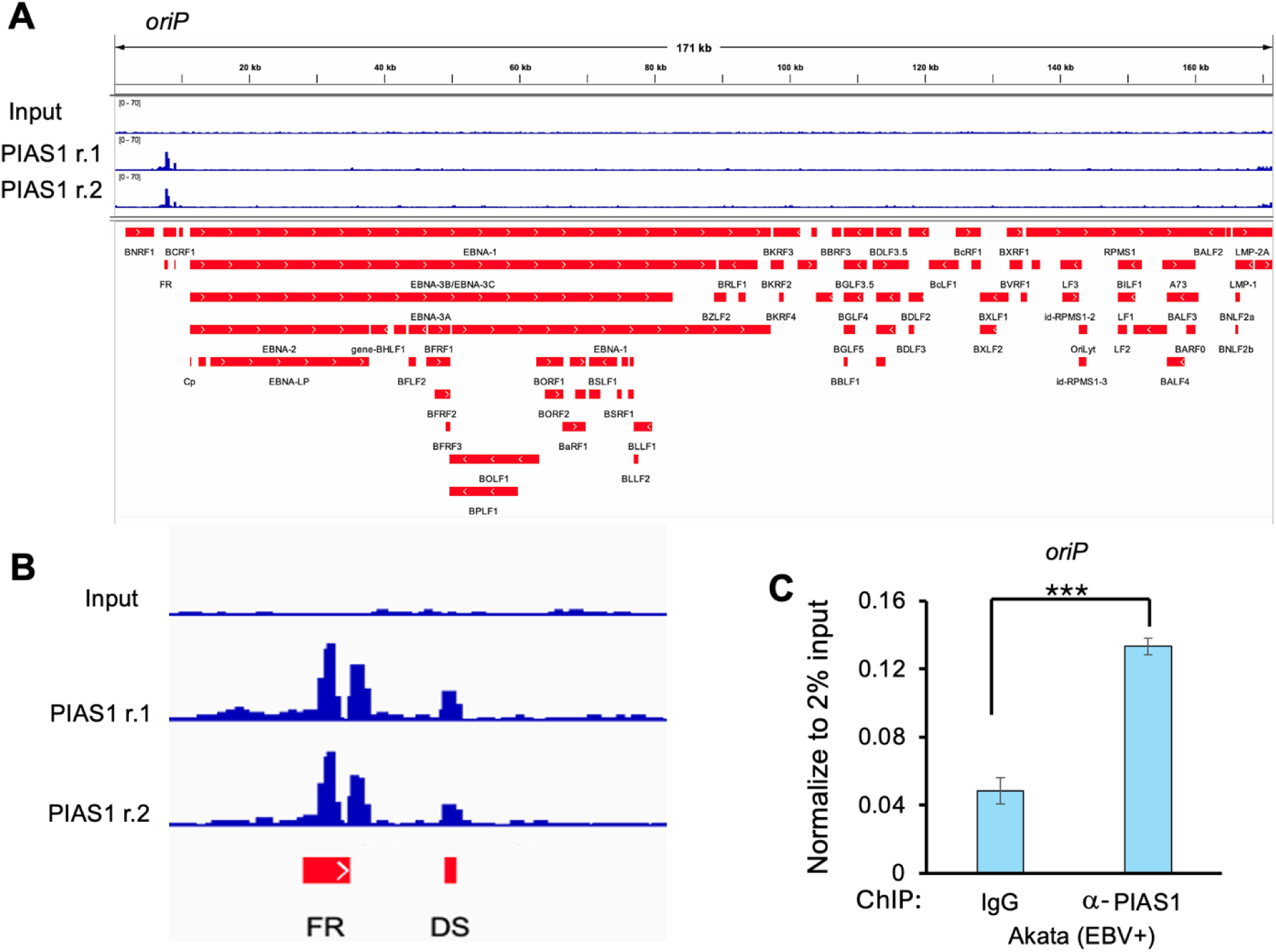
PIAS1 binds to EBV *oriP*. Akata (EBV+) cells were subjected to ChIP-Seq analysis to map PIAS1 DNA-binding regions across EBV genome. (**A**) Integrative Genomics Viewer (IGV) visualization of the EBV genome displaying ChIP-Seq tracks: Input (top), PIAS1 replicate 1 (PIAS1 r.1, middle), PIAS1 replicate 2 (PIAS1 r.2, bottom), along with EBV genome annotation (Akata strain). (**B**) Enlarged view highlighting PIAS1 binding at the *oriP* region of the EBV genome. FR, family of repeats; DS, dyad symmetry. (**C**) ChIP-qPCR analysis of the *oriP* region using anti-PIAS1 and control IgG antibodies according to the protocol outlined at Material and Methods. Data represents #x002B; SD from three biological replicates. ***p < 0.001.

### PIAS1 interacts with EBNA1

The EBV *oriP* region is known to be occupied by EBNA1, which is crucial for maintaining the EBV genome and facilitating replication [6, 7]. Based on PIAS1 ChIP-seq results (**Fig. 1**), we hypothesized that PIAS1 interacts with EBNA1 at the site of *oriP*. To test this hypothesis, we employed multiple experimental approaches in various cell lines. First, we transfected HEK-293T cells with plasmids expressing EBNA1 and PIAS1. Co-immunoprecipitation (co-IP) using anti-V5 antibody-conjugated magnetic beads, followed by Western blotting (WB) analysis with anti-PIAS1 antibody, demonstrated that PIAS1 co-immunoprecipitated with EBNA1 (**Fig. 2A**). To validate these findings in the context of EBV genome, we repeated the experiment in HEK-293 (EBV+) cells and obtained similar results (**Fig. 2B**).

To further confirm the interaction between PIAS1 and EBNA1 under physiological conditions, we conducted a proximity ligation assay (PLA) in Akata (EBV+) B cells. The cells were subjected to PLA with or without anti-PIAS1 and anti-EBNA1 antibodies. No PLA signals were detected in the absence of antibodies, whereas dot-like signals were observed in cells treated with both antibodies, primarily localized in the nucleus (**Fig. 2C**). To extend these findings to EBV+ epithelial cells, we repeated the PLA experiment in SNU-719, an EBV+ gastric cancer cell line. Consistent with those results in Akata (EBV+) cells, we observed strong PLA signals in SNU-719 cells using anti-PIAS1 and anti-EBNA1 antibodies (**Fig. 2D**). Collectively, these findings provide strong evidence for the interaction between PIAS1 and EBNA1 across multiple cell types, including B cells and epithelial cells.

**Figure 2.**
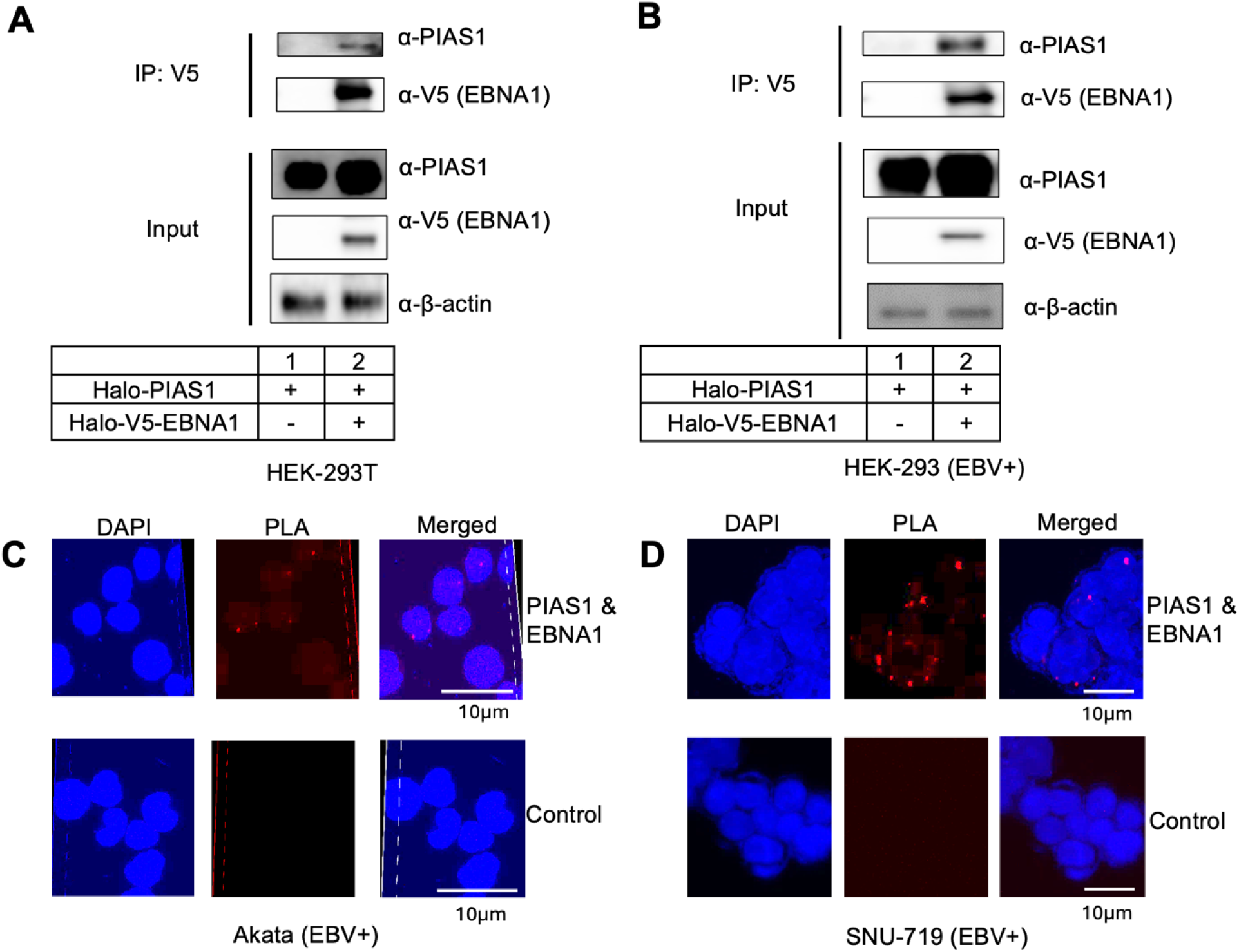
PIAS1 interacts with EBNA1. **(A-B)** HEK-293T (A) and HEK-293 (EBV+) (B) cells were co-transfected with PIAS1 and V5-EBNA1 as specified. WB analysis demonstrated that PIAS1 is co-IPed with EBNA1. Whole-cell lysates were probed for PIAS1 and V5-EBNA1 to confirm input levels. β-Actin was used as a loading control. (**C-D**) Akata (EBV+) (C) and SNU-719 (D) cells were blocked with 3% bovine serum albumin (BSA) in phosphate-buffered saline (PBS) for 1 hour at room temperature. Subsequently, the cells were incubated with either PBS control or a combination of mouse anti-EBNA1 and rabbit anti-PIAS1 antibodies. Probes were then added for ligation and amplification. Cell nuclei were visualized using Nikon AXR after staining with DAPI (4′,6-diamidino-2-phenylindole). The interaction between EBNA1 and PIAS1 *in situ* was indicated by red dots representing PLA signals.

To elucidate the specific regions of PIAS1 responsible for this interaction, we conducted co-IP experiments using HEK-293T cells transfected with plasmids expressing full-length HA-EBNA1 and either full-length or individual fragments of V5 tagged PIAS1 (**Fig. 3A**). The results revealed that full-length PIAS1 and all PIAS1 fragments except the C-terminal region (aa 409-651) are co-IPed with HA-EBNA1 (**Fig. 3B, lane 2, 3, and 5**). These observations suggest that the SAP and RING domains of PIAS1 are crucial for its interaction with EBNA1.

To identify the region(s) of EBNA1 involved in PIAS1 binding, we co-transfected HEK-293T cells with HA-EBNA1 fragments and V5-PIAS1 (**Fig. 3C**). Subsequent immunoprecipitation of PIAS1 using anti-V5 antibody-conjugated beads revealed that the C-terminal region (aa 216-405) of EBNA1, which contains the DNA binding and dimerization domain, is essential for PIAS1 interaction (**Fig. 3D, lane 6 vs lane 4**). Collectively, our findings indicate that the SAP and RING domains of PIAS1 interact specifically with the DNA binding domain of EBNA1 (**Fig. 3E**).

**Figure 3.**
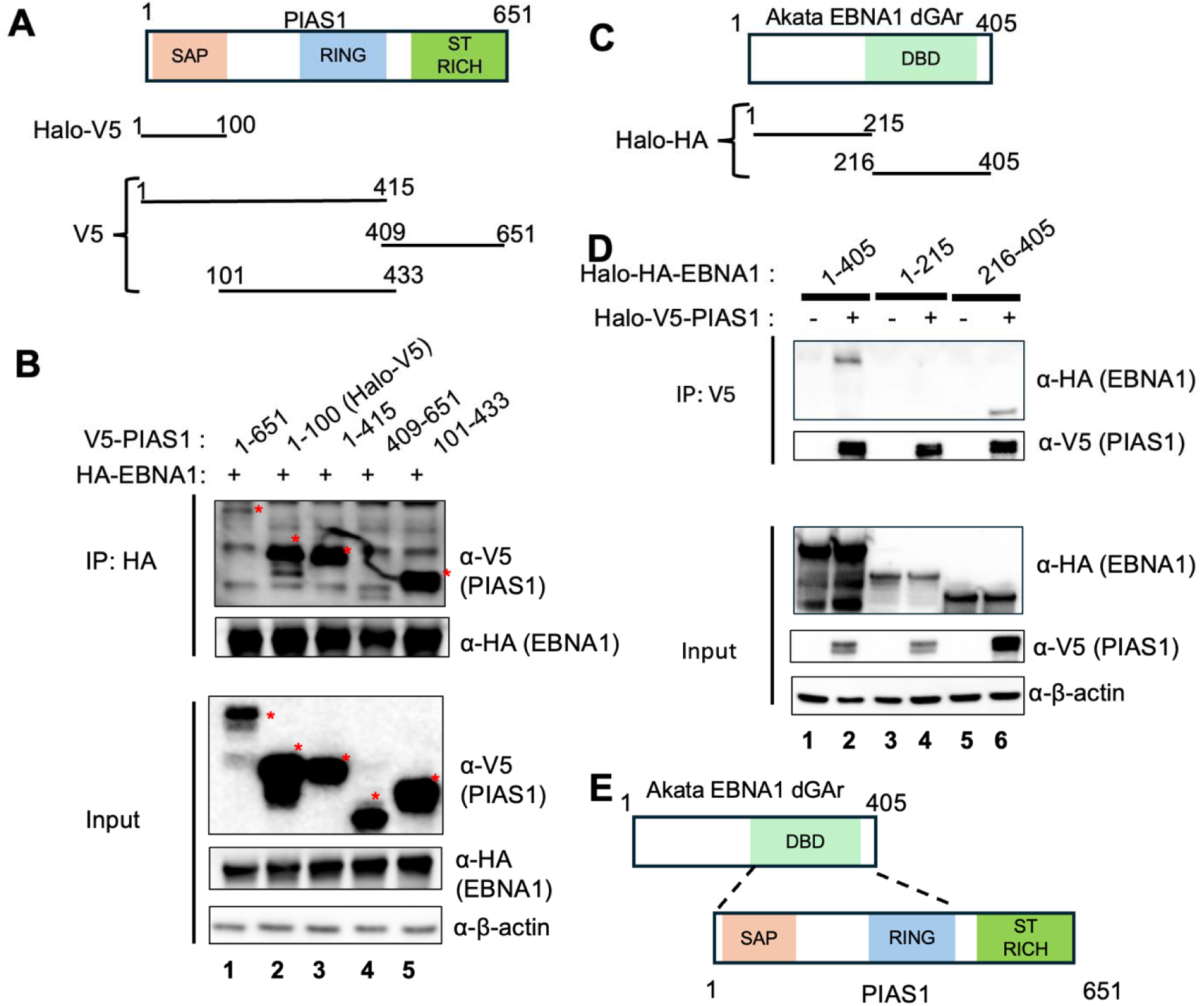
PIAS1 interacts with EBNA1 through their specific domains. (**A**) Schematic representation of PIAS1 domains and V5-or Halo-V5-PIAS1 truncation mutants. SAP, SAF-A/B, Acinus, and PIAS; RING, RING finger E3 ligase domain; ST-rich, Ser/Thr-rich region. (**B**) HEK-293T cells were co-transfected with EBNA1, full-length and truncated PIAS1 plasmids as indicated. WB analyses show that N-terminal and middle regions of PIAS1 are co-IPed with EBNA1. β-actin blot was included as a loading control. Asterisks denote the positions of full-length and truncated PIAS1. (**C**) Schematic representation of EBNA1 domains and Halo-V5-EBNA1 truncation mutants. DBD, DNA binding domain. (**D**) HEK-293T cells were co-transfected with full-length Halo-V5-PIAS1 and full-length or truncated Halo-HA-EBNA1 as indicated. WB analyses show that C-terminal region of EBNA1 is co-IPed with PIAS1. β-actin blot was included as a loading control. (E) A proposed model showing the C-terminal regions of EBNA1 binding to the N-terminal and middle regions of PIAS1.

### PIAS1 enhances EBNA1 SUMOylation in both *in vivo* and *in vitro*

Our previous studies have demonstrated that PIAS1 enhances the anti-viral activity of SAMHD1 and YTHDF2 by promoting their SUMOylation on multiple lysine residues [18, 19]. The E3 ligase responsible for EBNA1 SUMOylation remained unidentified, despite a previous report confirming EBNA1 as a SUMOylated protein [23]. We hypothesized that PIAS1 functions as the E3 ligase facilitating EBNA1 SUMOylation. To test this hypothesis, we transfected HEK-293T cells with plasmids expressing EBNA1, PIAS1, and SUMO2 (**Fig. 4A**). Our results showed that while individual transfection of PIAS1 slightly increased total SUMOylation levels, co-transfection of SUMO2 and PIAS1 significantly enhanced SUMOylation (**Fig. 4A lane 4 vs lane 3**).

To further investigate PIAS1-mediated SUMOylation of EBNA1, we performed IP using anti-V5 antibody-conjugated magnetic beads. WB analysis with anti-SUMO2/SUMO3 antibody revealed a strong SUMOylated EBNA1 band when SUMO2 and PIAS1 were co-expressed, suggesting that EBNA1 is indeed targeted for SUMOylation by PIAS1 (**Fig. 4 lane 8**).

The robust interaction between PIAS1 and EBNA1, coupled with the strong SUMOylated EBNA1 signal when PIAS1 is present, we reasoned that PIAS1 directly SUMOylates EBNA1. To examine this further, we conducted *in vitro* SUMOylation experiments using purified V5-EBNA1 and purified PIAS1 in combination with E1, E2 (UBC9) and SUMO2. Our findings showed that PIAS1 also enhances SUMOylation of EBNA1 *in vitro* (**Fig. 4B lane 3 vs lane 1 and 2**), suggesting EBNA1 is a direct substrate for PIAS1.

**Figure 4.**
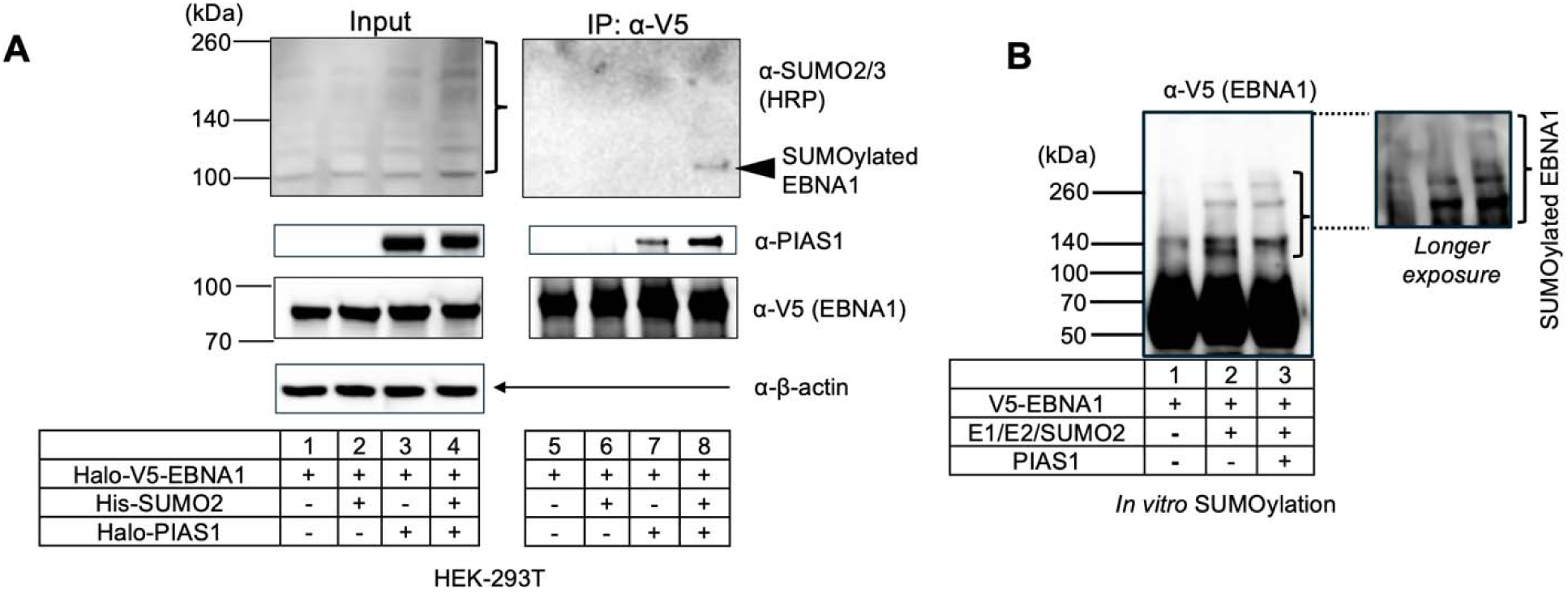
PIAS1 enhances EBNA1 SUMOylation both *in vivo* and *in vitro*. (**A**) HEK-293T cells were transfected with plasmids encoding Halo-V5-EBNA1, Halo-PIAS1, and His-SUMO2. Whole-cell lysates (input) were analyzed by WB using antibodies against SUMO2/3, PIAS1, V5, and β-actin. SUMOylated proteins are indicated by brackets. EBNA1 was immunoprecipitated using anti-V5 magnetic beads, followed by WB analysis with antibodies as indicated. Arrows denote SUMOylated EBNA1. IP denote immunoprecipitation. (**B**) *In vitro* SUMOylation assay was conducted using a combination of purified proteins, including E1, E2, SUMO2, PIAS1, and the substrate V5-EBNA1, as specified. The reaction wa stopped by adding 2X SDS-PAGE loading buffer, followed by WB analysis using anti-V5-HRP antibody. SUMOylated EBNA1 is indicated by brackets.

### PIAS1 mediates SUMOylation of EBNA1 at three lysine residues

SUMOylation typically occurs on lysine residues within the consensus motif ΨKxE/D or the inverted motif E/DxKΨ (Ψ represents a hydrophobic amino acid, x can be any amino acid). However, SUMOylation can also occur outside these consensus sequences [24]. To identify SUMOylation sites on EBNA1, we first used GPS-SUMO 2.0 webserver (https://sumo.biocuckoo.cn/advanced.php) to predict putative SUMOylation sites.

The program identified three high-scoring SUMOylation sites: K241, K17, and K75. K241 (P**K**FE) and K17 (Q**K**ED) are located within the KxE/D motif but motif in K17 lacks a hydrophobic amino acid, while K75 (Q**K**RP) belongs to a non-consensus motif (**Fig. 5A**). To verify PIAS1-mediated SUMOylation at these sites, we created individual lysine-to-arginine mutants and performed *in vitro* SUMOylation assays, including pCEP4 as an EBV replicon containing *oriP* to mimic EBNA1 binding conditions.

Although K241 was previously reported as the major SUMOylation site, our *in vitro* assays showed substantial SUMOylation signals for each individual mutant (**Fig. S1, lanes 4, 6, 8 vs. lane 2**). We then generated a triple mutant, K17R/K75R/K241R (RRR), which greatly reduced SUMOylation signal compared to the wild-type protein in the *in vitro* assay (**Fig. 5B lane 8 vs lane 4**). The addition of *oriP* plasmid stimulated EBNA1 SUMOylation by PIAS1, suggesting *oriP* binding enhances the interaction between PIAS1 and EBNA1 (**Fig. 5B lane 3 vs lane 4**). These results demonstrate that K17 (N-terminal), K75 (N-terminal), and K241 (C-terminal) are the 3 major SUMOylation sites on EBNA1 mediated by PIAS1, with individual sites capable of compensating for the loss of others (**Fig. 5C**).

According to the AlphaFold3-predicted three-dimensional structure of EBNA1 [25], lysine residues K17 and K75 are situated within an intrinsically disordered region, whereas K241 resides within an α-helix region. The localization of lysines within or adjacent to these flexible regiones should enhance structural accessibility, thereby facilitating SUMOylation (**Fig. 5D**).

To investigate the conservation of EBNA1 SUMOylation sites across various EBV strains and the closed related species, we aligned the amino acid sequences of six EBV EBNA1 with two cyno-EBV EBNA1 sequences from viruses that infect cynomolgus macaques [26]. Notably, we found that the amino acids corresponding to K75/K241 of Akata EBNA1 are conserved among all examined species. However, K17 of Akata EBNA1 are conserved only in EBV strains but absent in cyno-EBV EBNA1 (K to R). This observation suggests EBNA1 SUMOylation by PIAS1 is likely conserved in different EBV and cyno-EBV strains (**Fig. 5E**).

**Figure 5.**
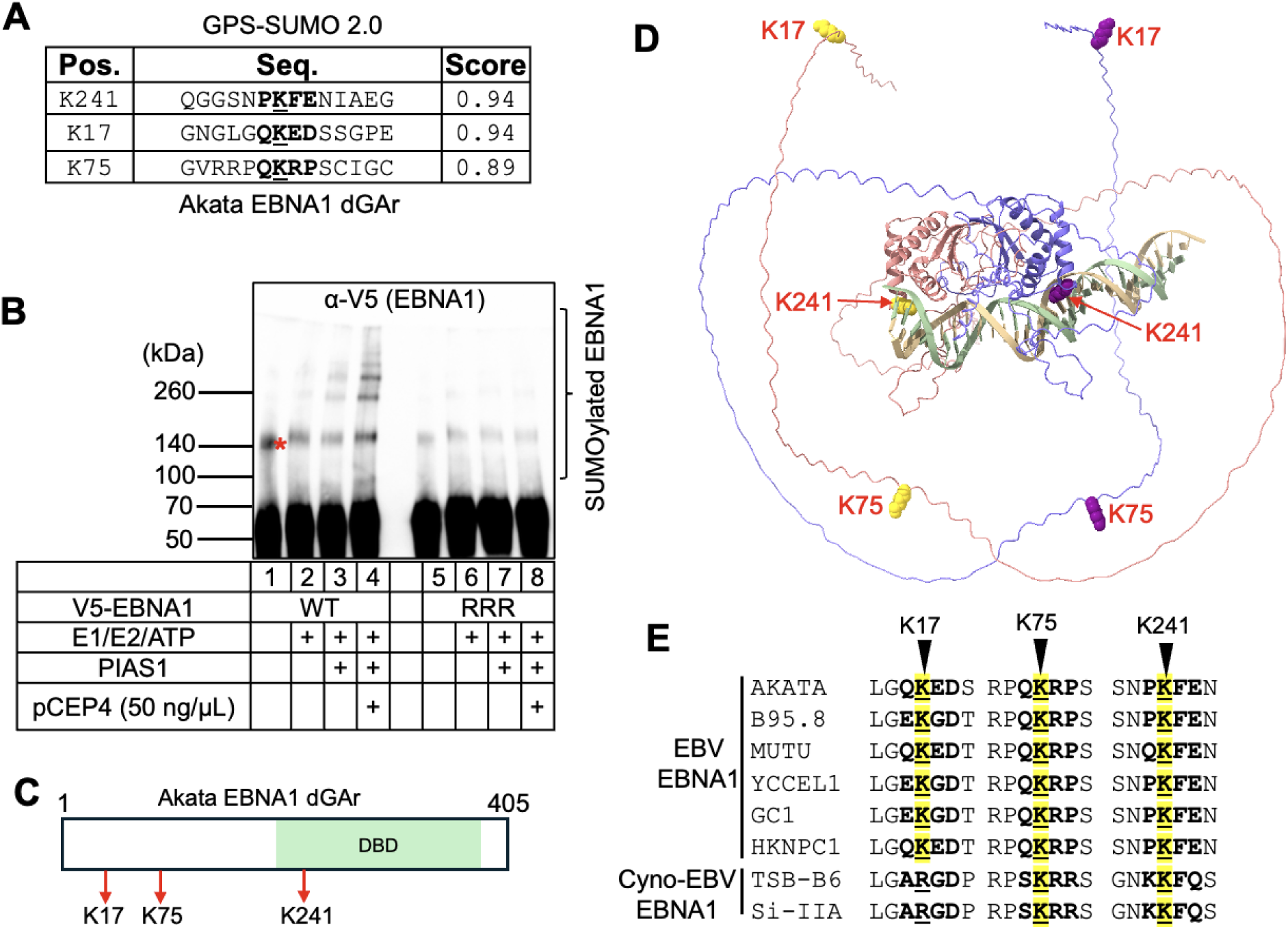
PIAS1 mediates SUMOylation of EBNA1 at three lysine residues. (**A**) SUMOylation sites on EBNA1 were predicted using the GPS-SUMO 2.0 webserver. (**B**) *In vitro* SUMOylation assay was performed on wild-type EBNA1 and a triple lysine-to-arginine mutant (K17R/K75R/K241R; RRR mutant) using E1, E2, SUMO2, and *oriP*-containing plasmid (pCEP4) as indicated. WB analysis was performed with anti-V5-HRP antibody after terminating the reaction with 2X SDS-PAGE loading buffer. Red asteris denotes non-specific band; Bracket denotes SUMOylated EBNA1. (**C**) A schematic representation of the EBNA1 (dGAr) protein structure highlights the relevant lysine residues with red arrows. (**D**) The predicted three-dimensional structure of EBNA1 (dGAr) dimer with 1XFR DNA sequence (5’-GGATAGCATATACTACCCGGATATAGATTA-3’), generated using AlphaFold3, shows the three SUMOylation sites (K17, K75, and K241). (**E**) Sequence alignment of EBNA1 from 6 EBV strains and 2 cynomolgus macaque EBV (cyno-EBV) using CLUSTAL OMEGA [27]. The conserved SUMOylation sites were highlighted in yellow.

**Figure S1.**
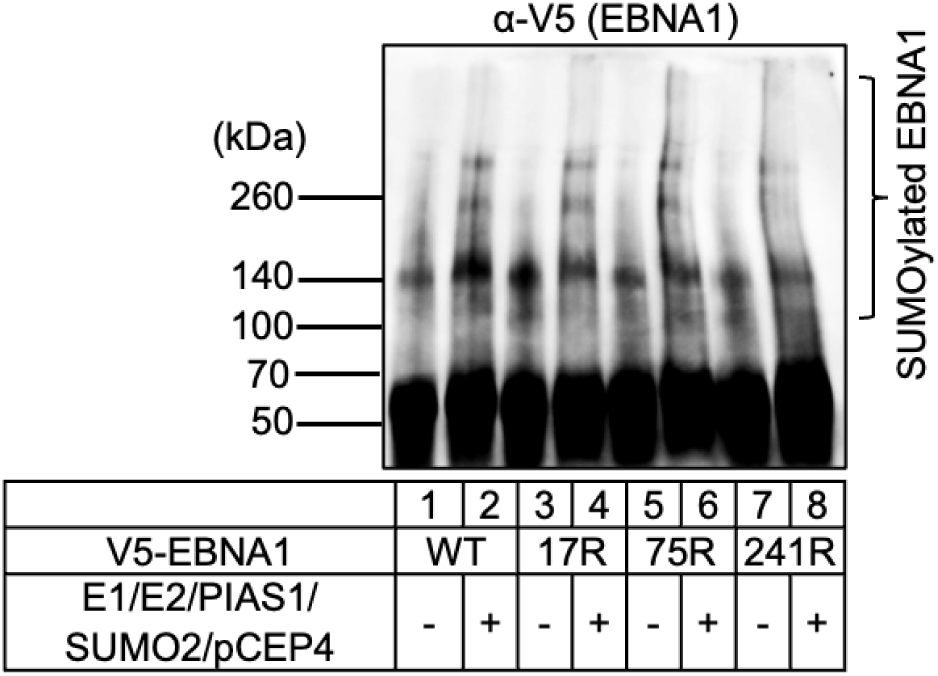
The impact of K17R, K75R and K241R individual mutations on EBNA1 SUMOylation by PIAS1. Wild-type (WT) EBNA1 and single-site mutants (K17R, K75R, and K241R) were subjected to SUMOylation using E1, E2, SUMO2, PIAS1 and *oriP*-containing plasmid pCEP4. WB analysis was performed with anti-V5-HRP antibody. Bracket denotes poly-SUMOylated protein.

To investigate whether SUMOylation site mutations affect EBNA1 dimerization, we co-transfected WT HA-EBNA1 and V5-EBNA1 (WT or SUMO-deficient mutants) into HEK-293 (EBV+) cells and performed co-immunoprecipitation using anti-V5 magnetic beads. Western blot analysis of HA-EBNA1 showed no significant difference in dimerization between WT and SUMOylation-deficient EBNA1 (**Fig. 6A**). We also assessed DNA binding affinity of EBNA1 WT and the RRR mutant using Electrophoretic Mobility Shift Assays (EMSA) with increasing protein concentrations (0–500 nM) and a 2X FR probe (10 nM). The EMSA results revealed no detectable difference in DNA-binding capacity between WT EBNA1 and RRR mutant (**Fig. 6B**). To examine whether PIAS1 binds to *oriP* DNA directly, we also performed EMSA using PIAS1 protein and a 2X FR probe. We found that there is weak EMSA signals only at higher concentrations of PIAS1 (**Fig. 6C**).

To evaluate whether PIAS1 affects EBNA1’s DNA binding capability, we then performed EMSA with a fixed concentration of EBNA1 and increasing amounts of PIAS1. We found that with increasing PIAS1 concentrations, there are a gradual decrease of free probes and a trace increase of EMSA signals (**Fig. 6D**), suggesting PIAS1 and EBNA1 may coordinate to binds to *oriP* DNA.

**Figure 6.**
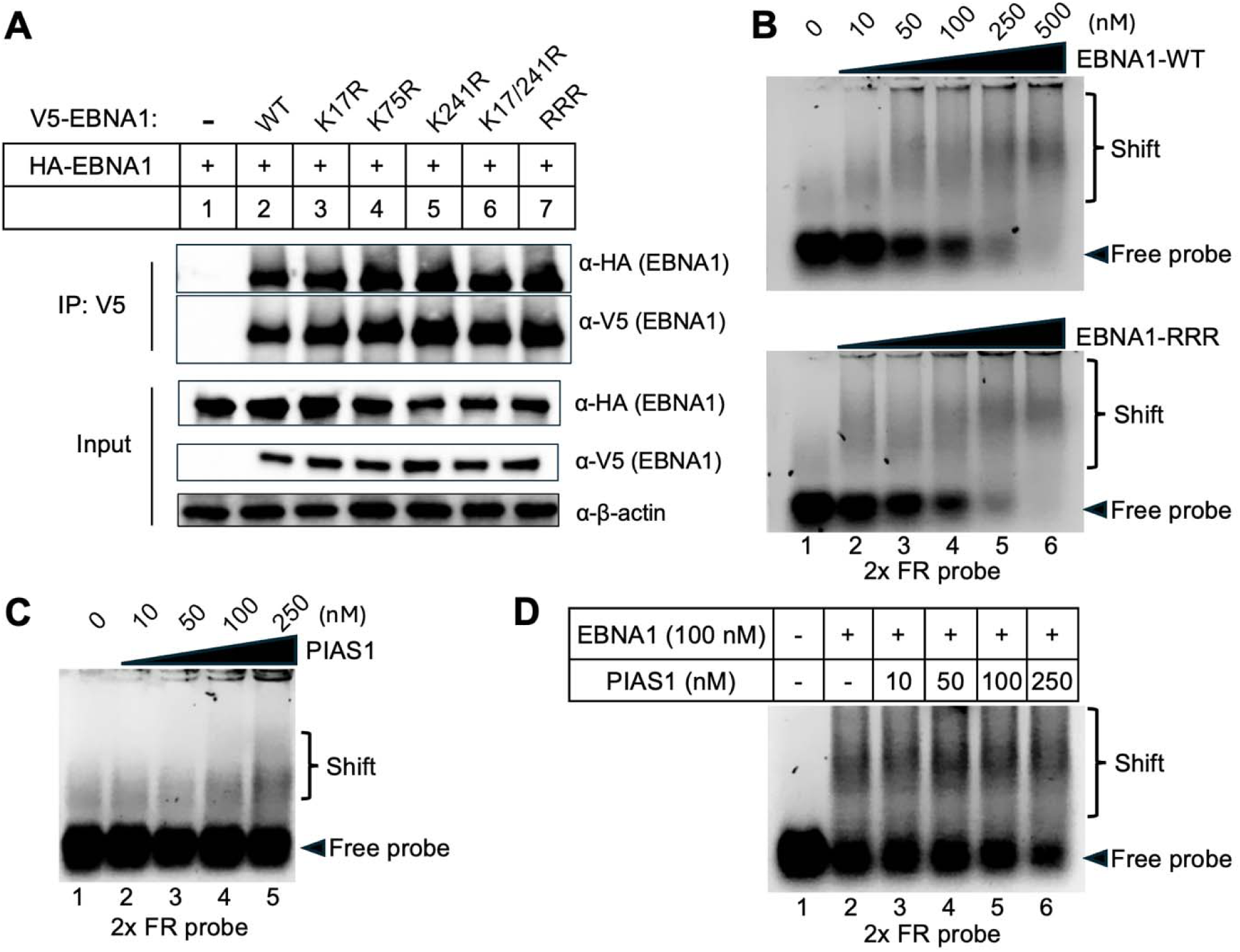
SUMOylation-deficient EBNA1 maintains dimerization and DNA binding capabilities. (**A**) HEK-293 (EBV+) cells were co-transfected with HA-EBNA1 WT and V5-EBNA1 (WT or mutants as indicated). WB analysis demonstrates that HA-EBNA1 is co-IPed with WT and mutant V5-EBNA1. Input: whole-cell lysates probed for HA-EBNA1, V5-EBNA1, and β-actin (loading control). (**B**) EMSA was performed using increasing concentrations of purified EBNA1 (WT and RRR mutant) and a 2X FR probe. The shifted and free probes were resolved on 1.4% agarose gel in 1x TBE buffer. (**C**) EMSA was performed using increasing concentrations of purified PIAS1 and a 2X FR probe. (**D**) EMSA was performed using a fixed concentration of EBNA1, increasing concentrations of PIAS1 and a 2X FR probe.

**Figure 7.**
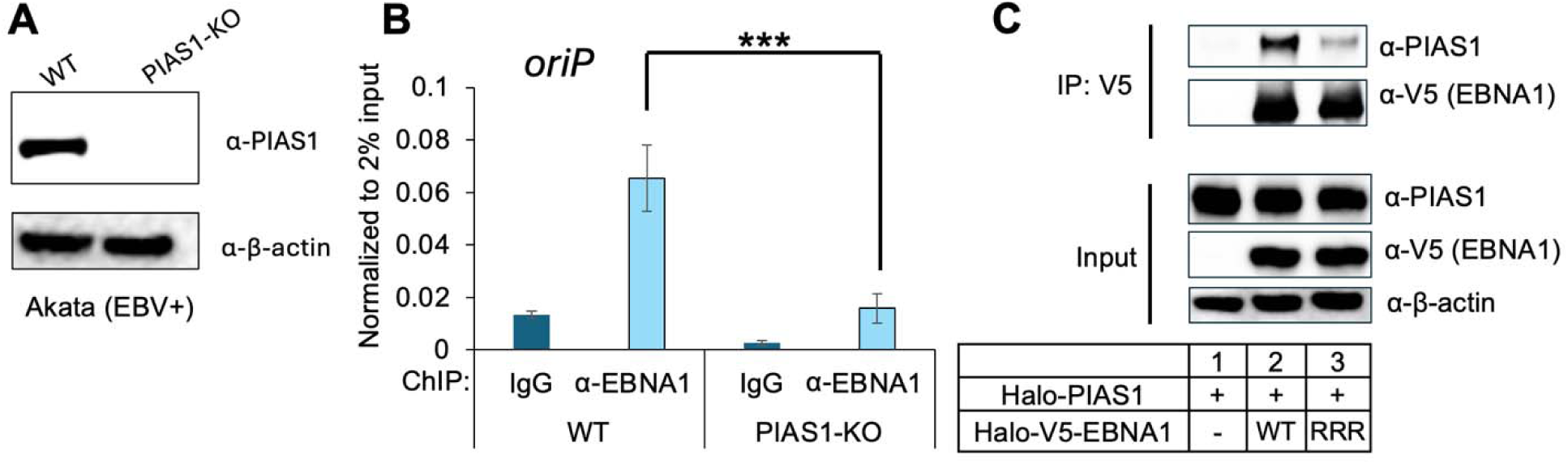
PIAS1 promotes EBNA1 binding to *oriP*. (**A**) WB analysis of PIAS1 and β-actin expression levels between wild-type (WT) and PIAS1-knockout (PIAS1-KO) Akata (EBV+) cells. (**B**) ChIP-PCR analysis of EBNA1 binding to *oriP* in WT and PIAS1-KO Akata (EBV+) cells. Nonspecific IgG serves as a negative control. Data represents SD from three biological replicates. ***p < 0.001.(**C**) HEK-293T cells were co-transfected with PIAS1 and V5-EBNA1 (WT or RRR mutant). WB analysis shows PIAS1 is co-IPed with EBNA1. Input: whole-cell lysates blotted for PIAS1, V5-EBNA1, and β-actin (loading control).

To determine whether PIAS1 affects EBNA1 binding to *oriP*, we performed ChIP analysis for EBNA1 in WT and PIAS1-knockout (PIAS1-KO) Akata (EBV+) cells (**Fig. 7A**). Interestingly, we found that PIAS1-KO significantly reduces EBNA1 binding to *oriP* (**Fig. 7B**). SUMOylation of substrate also could in turn affect its interaction with PIAS1, as PIAS1 has multiple SUMO-interacting motifs [28]. To examine whether SUMOylation-deficient EBNA1 affects its binding with PIAS1, we performed co-IP experiment comparing PIAS1 binding to WT and RRR mutant EBNA1. We noticed that PIAS1 binding to EBNA1-RRR mutant is significantly reduced compared to WT EBNA1 (**Fig. 7C, lane 3 vs lane 2**), suggesting that SUMOylation of EBNA1 promotes its interaction with PIAS1.

**Figure 8.**
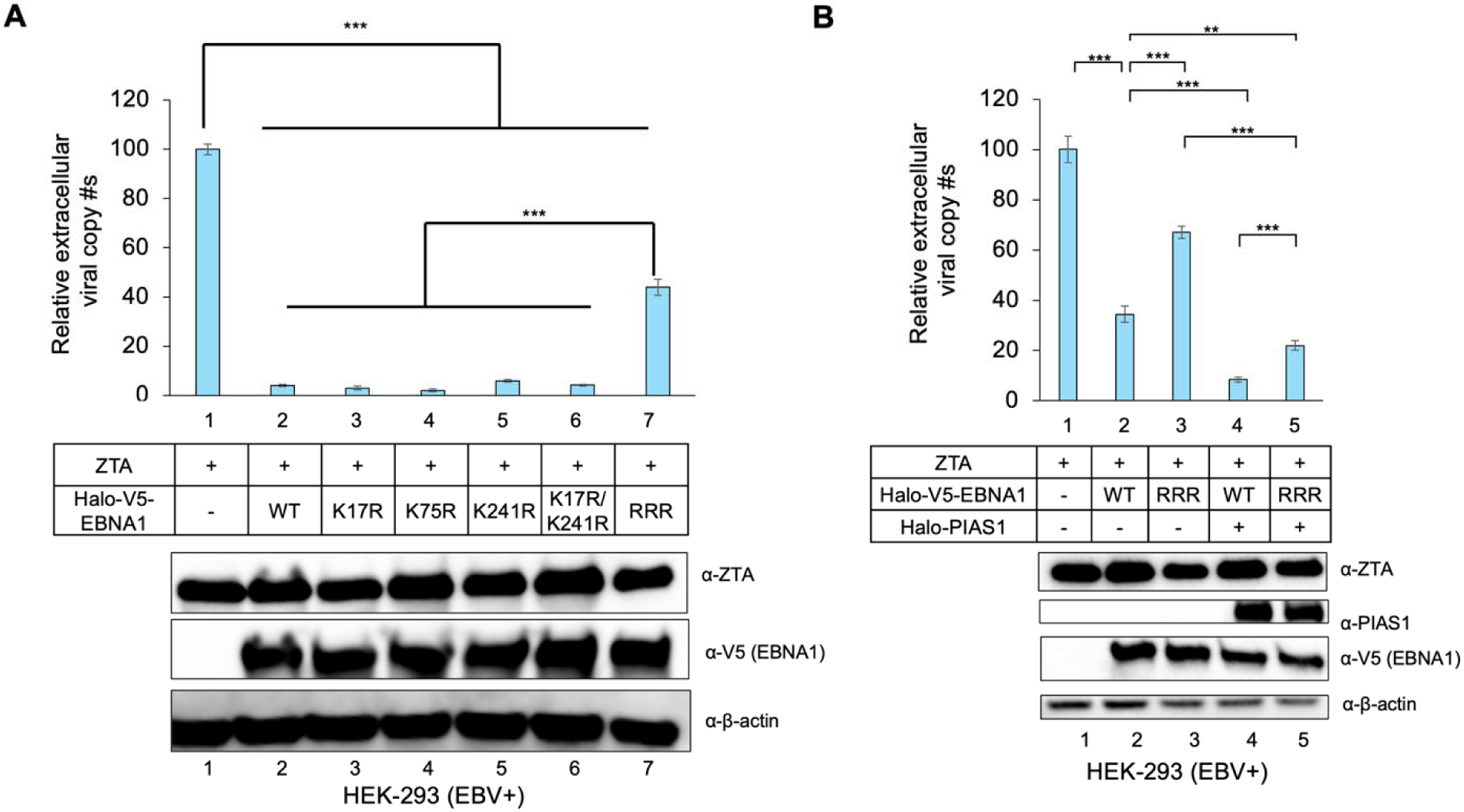
PIAS1 synergizes with EBNA1 to repress EBV lytic replication. (**A**) SUMOylation-deficient EBNA1 impairs its ability to restrict EBV replication. HEK-293 (EBV+) cells were co-transfected with ZTA plasmid (lytic inducer), and either WT EBNA1 or EBNA1 mutants as indicated. Relative EBV cop numbers were quantified using qPCR according to the protocol outlined in the Materials and Method section. The value in lane 1 was normalized to 100. WB was employed to monitor the expression levels of ZTA and V5-EBNA1, with β-actin serving as a loading control. Data represents SD from three biological replicates. ***p < 0.001. (**B**) PIAS1 and EBNA1 synergistically repress EBV lytic replication. HEK-293 (EBV+) cells were co-transfected with ZTA plasmid (lytic inducer), PIAS1, and either wild-type WT EBNA1 or SUMOylation-deficient EBNA1 (RRR). Relative EBV copy numbers were quantified using qPCR. The value in lane 1 was normalized to 100. WB was employed to monitor the expression levels of ZTA, PIAS1, and V5-EBNA1, with β-actin serving as a loading control. Data represents SD from three biological replicates. **p < 0.01; ***p < 0.001

### PIAS1 synergizes with EBNA1 to repress EBV lytic replication

As previously reported, EBNA1 expression inhibits spontaneous EBV reactivation [29]. To investigate the role of EBNA1 SUMOylation in EBV replication, we compared WT EBNA1 with various EBNA1 SUMOylation site mutants using HEK-293 (EBV+) cells as a model. We found that SUMOylation-deficient RRR mutant was impaired to restrict EBV replication compared to WT EBNA1, individual K to R mutants, and K17R/K241R double mutant (**Fig. 8A lane 7 vs lane 2-6**).

To examine whether PIAS1 synergizes with EBNA1 to restrict EBV replication, we co-transfected HEK-293 (EBV+) cells with PIAS1, WT EBNA1, and the RRR mutant. Without PIAS1, the RRR mutant showed increased EBV lytic replication compared to WT EBNA1 (**Fig. 8B lane 3 vs lane 2**). In the presence of PIAS1, both WT EBNA1 and RRR mutant further inhibited EBV replication, with WT EBNA1 having the strongest effect (**Fig. 8B lane 4-5 vs lane 2-3**). These results together suggested that both EBNA1 SUMOylation and its interaction with PIAS1 contribute to reduced EBV lytic replication.

**Figure 9.**
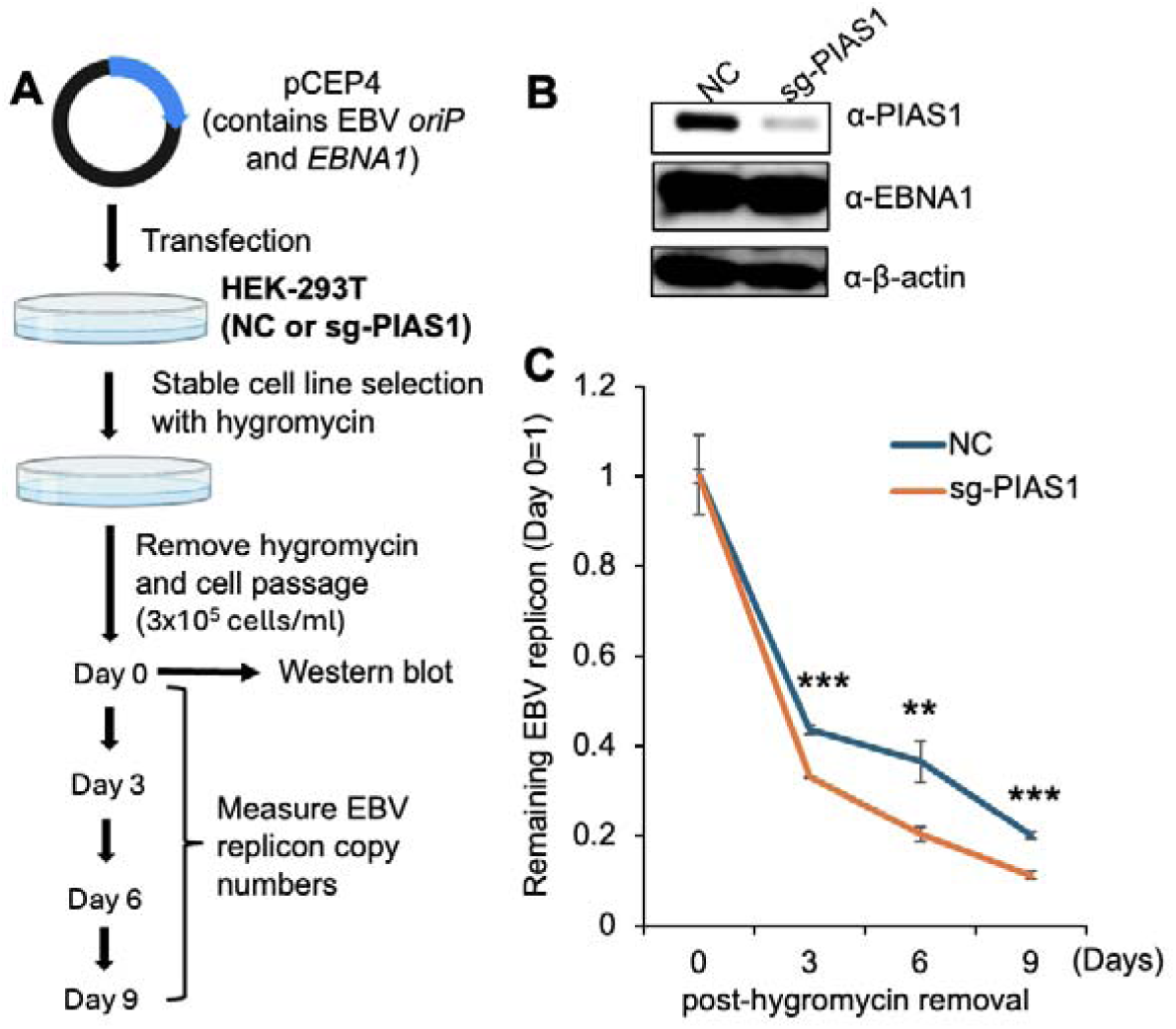
Loss of PIAS1 accelerates EBV replicon loss. (**A**) Schematic representation of EBV replicon (pCEP4-based plasmid) retention assays using HEK-293T cells carrying non-targeting control sgRNA (NC) and PIAS1-targeting sgRNA (sg-PIAS1). (**B**) WB analysis comparing PIAS1, EBNA1, and β-actin expression in HEK-293T cells carrying NC and sg-PIAS1. (**C**) The remaining of EBV replicon was measured by qPCR over 9 days after removal of hygromycin B. Data represents SD from three biological replicates. ** p<0.01; ***p < 0.001.

### PIAS1 promotes the maintenance of *oriP*-based plasmid by EBNA1 through SUMOylation

EBNA1 is known to function in EBV genome maintenance. To investigate the role of SUMOylation in this process, we utilized pCEP4 plasmid as an EBV *oriP*-based replicon that expresses EBNA1 and contains *oriP* for plasmid maintenance. We established pCEP4 stable cell line in HEK-293T cells carrying non-targeting control sgRNA (NC) or PIAS1 targeting sgRNA (sg-PIAS1) cells under hygromycin B selection (**Fig. 9A-B**). To study plasmid maintenance, we conducted serial passages every 3 days without hygromycin B to observe plasmid maintenance rates. We observed faster plasmid loss in PIAS1-depleted cells compared to control cells over 9 days (**Fig. 9C**), suggesting that PIAS1 contributes to pCEP4 plasmid retention.

**Figure 10.**
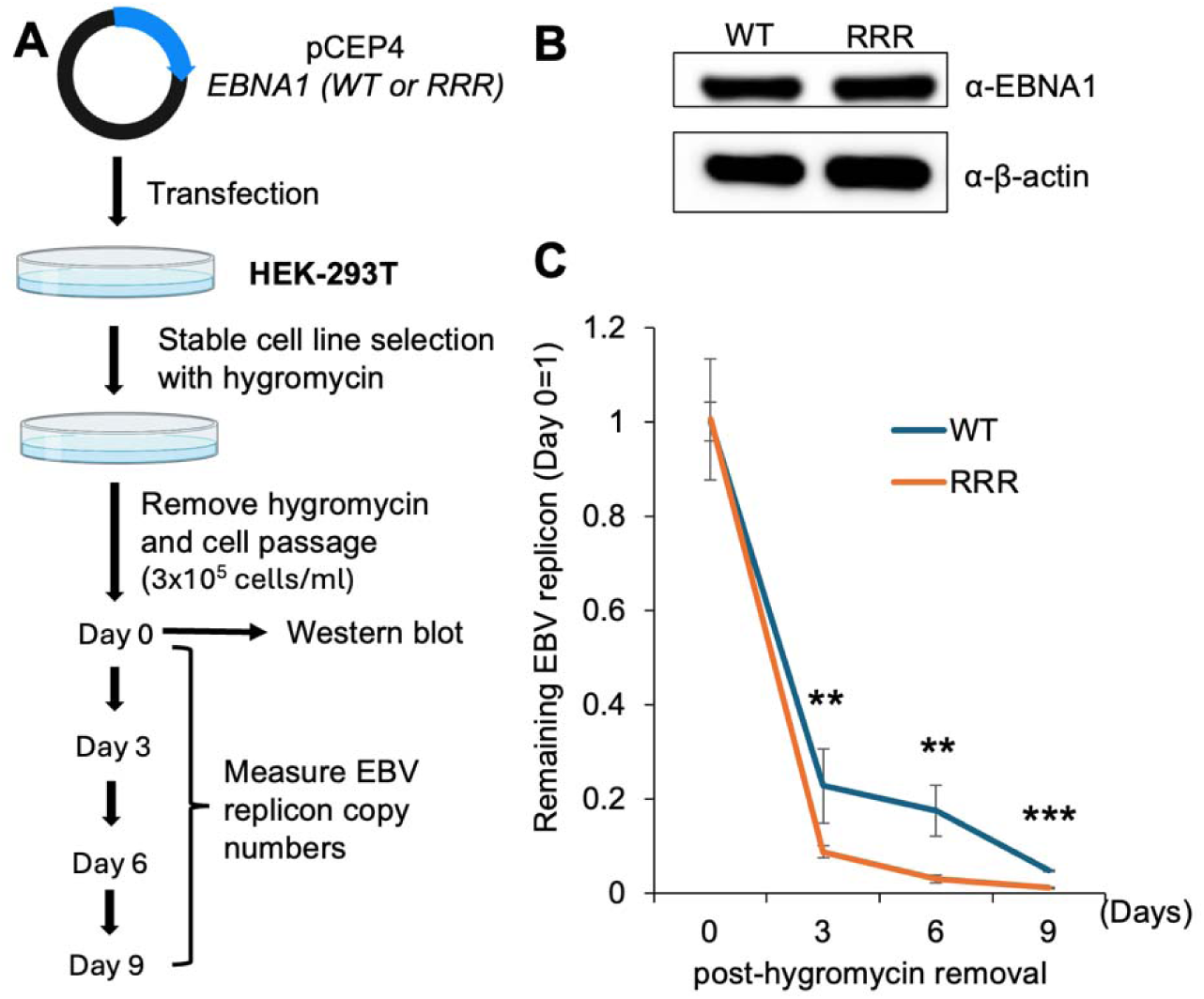
SUMOylation-deficient EBNA1 facilitates EBV replicon loss. (**A**) Schematic representation of EBV replicon retention assays using HEK-293T cells carrying WT EBNA1 and RRR mutant plasmids. (**B**) WB analysis comparing EBNA1 and β-actin expression in HEK-293T cells carrying pCEP4-EBNA1-WT and pCEP4-EBNA1-RRR plasmids (**C**) The remaining of EBV replicon was measured by qPCR over 9 days after removal of hygromycin B. Data represents ±SD from three biological replicates. ** p<0.01; ***p < 0.001.

To investigate the direct impact of EBNA1 SUMOylation on plasmid maintenance, we then mutated EBNA1 within pCEP4 to generate SUMOylation-deficient EBNA1 (RRR, K17R/K75R/K289R). The K289 residue corresponds to K241 in our EBNA1 dGAr due to additional GA repeats in pCEP4 plasmids.

We then transfected HEK-293T cells with both pCEP4 EBNA1-WT and the pCEP4 EBNA1-RRR plasmids, followed by hygromycin B selection (**Fig. 10A**). After establishing stable cell lines (**Fig. 10B**), we conducted plasmid retention assays in the absence of hygromycin B. We found that there is an increased loss of pCEP4 EBNA1-RRR plasmid compared to pCEP4 EBNA1-WT plasmid over 9 days (**Fig. 10C**), indicating that EBNA1 SUMOylation promotes *oriP*-based plasmid retention.

## DISCUSSION

EBNA1 is an essential EBV protein that plays key role in viral genome replication, episome maintenance, and transcriptional regulation. It is the only viral protein consistently expressed across all types of EBV latency and in EBV-associated malignancies. Structurally, EBNA1 contains a GAr region, DNA-binding and dimerization domains, and nuclear localization signals. Its primary functions include tethering the viral episome to host chromosomes during cell division and modulating the expression of both viral and host genes [3, 5, 6, 8].

It was reported that EBNA1 is regulated by various post-translational modifications (PTMs), such as phosphorylation, arginine methylation, lysine hydroxylation, and SUMOylation. Phosphorylation is a key modification for EBNA1, with ten phosphorylated residues identified by mass spectrometry. These phosphorylation sites are located in the N-terminal GAr, glycine/arginine-rich domains (GR1 and GR2), and near the nuclear localization sequence. Phosphorylation-deficient mutants show reduced *oriP*-dependent transcription and episome maintenance, while retaining normal half-life and nuclear localization, highlighting the importance of this modification [30–32].

In addition, phosphorylation of EBNA1 at Ser393 by viral and cellular kinases may influence its antigenicity, modulate antibody response, and promote cross-reactivity with GlialCAM, a phenomenon observed in clonally expanded B cells in multiple sclerosis[33–35]

Arginine methylation, catalyzed by protein arginine methyltransferases (PRMTs), is another crucial PTM affecting EBNA1 stability, protein interactions, transcription activation, and episome maintenance. GAr region is particularly important for segregation and transcriptional activation functions [31, 36]. EBNA1 stability and DNA replication activity are regulated by PLOD1-mediated lysine hydroxylation, as well as ubiquitin-proteasome-dependent degradation [37, 38].

SUMOylation, the covalent attachment of small ubiquitin-like modifier (SUMO) proteins, has been implicated in the regulation of EBNA1’s functions. Previous studies suggested that loss of EBNA1 SUMOylation at K477 impairs viral DNA persistence and enhances spontaneous EBV reactivation [23]. It was reported that SUMO2 is covalently attached to K1140 of the KSHV-LANA, promoting viral genome maintenance and repressing lytic reactivation through inhibition of RTA expression [39]. In the case of HPV16, another study identified K292 of the E2 protein as a SUMOylation site, although its functional consequences remain unexplored [40].

In this study, we demonstrated that PIAS1 is specifically enriched at EBV *oriP* (**Fig. 1**), where it binds to and colocalizes with EBNA1 (**Fig. 2**). We further demonstrated that these interactions are mediated by the N-terminal and central regions of PIAS1 and the C-terminal DBD of EBNA1 (**Fig. 3**).

Importantly, we discovered that PIAS1 serves as an E3 SUMO ligase for EBNA1 (**Fig. 4**). In addition to the previously characterized K477 (corresponding to K241 in GAr deleted EBNA1), we identified two novel SUMOylation sites in EBNA1, namely K17 and K75 (**Fig. 5**). These two sites reside in non-consensus SUMOylation motifs. Mutation of individual lysine residue to arginine did not affect EBNA1 SUMOylation. However, the mutation of three sites abolished EBNA1 SUMOylation signals. These findings underscore the functional relevance and prevalence of non-consensus SUMOylation sites in EBNA1 regulation within cells (**Fig. 5**).

The binding of PIAS1 to EBNA1 was also implicated in EBNA1’s DNA binding where PIAS1 slightly promotes EBNA1 binding to *oriP* DNA (**Fig. 6**). The loss of PIAS1 also diminished EBNA1 binding to to *oriP* in Akata (EBV+) cells (**Fig. 7**), suggesting that PIAS1 plays a critical role for EBNA1’s chromatin binding activity at *oriP*.

As previously reported, both PIAS1 and EBNA1 have been implicated in EBV lytic reactivation [20, 29]. Intriguingly, we found that SUMOylation-deficient EBNA1 is compromised in limiting EBV lytic replication, and that PIAS1 synergizes with EBNA1 to further restict viral replication (**Fig. 8**). These findings expand the function of PIAS1-mediated SUMOylation from our previously identified targets, SAMHD1 and YTHDF2 [18, 19], to now include EBV EBNA1.

EBNA1 was recently reported to coordinate with H2A.Z for epigenetic reprogramming of EBV episomes [41]. Interestingly, we demonstrated that both PIAS1 binding and SUMOylation of EBNA1 contributes to the retention of EBV *oriP*-based replicon (**Figs. 9 and 10**), suggesting an important regulatory mechanism for EBV episome maintenance.

Our study paves the way for exploring whether KSHV LANA [39] and HPV E2 [40] are regulated by PIAS1. In co-transfection systems, we observed that PIAS1 interacts with KSHV LANA and HPV-16 E2, promoting their SUMOylation *in vitro* (**Fig. S2**). The functional significance of these interactions warrants further investigation, particularly in light of previous proteomic screens that identified PIAS1 as a potential E2-interacting partner [42].

**Figure S2.**
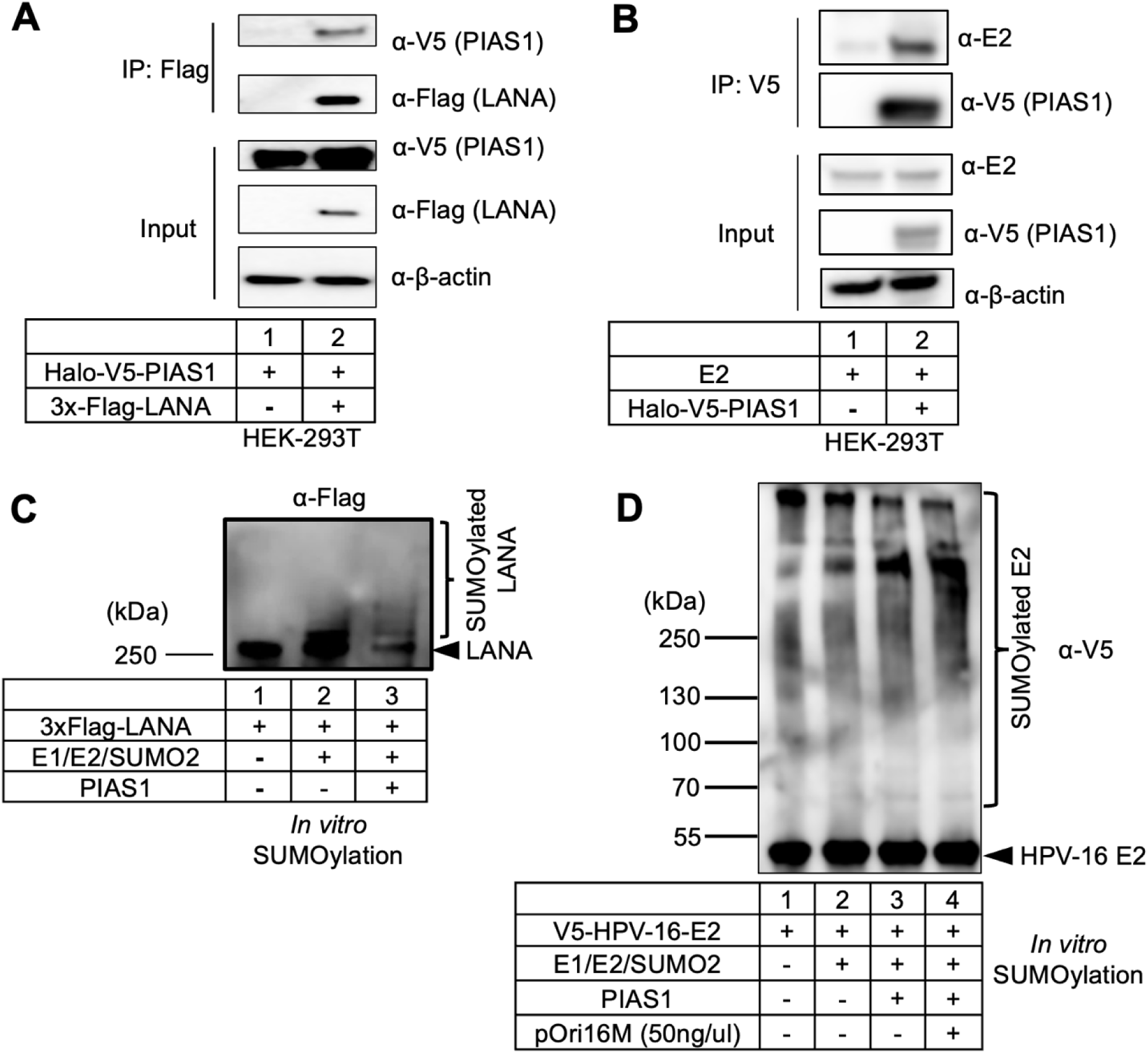
PIAS1 promotes the SUMOylation of KSHV-LANA and HPV-16 E2. (**A**) HEK-293T cells were co-transfected with V5-PIAS1 and 3x-Flag-LANA plasmids. WB analysis showing that V5-PIAS1 is co-IPed with 3x-FLAG-LANA. Input: whole-cell lysates probed for V5-PIAS1, Flag-LANA, and β-actin (loading control). (B) HEK-293T cells were co-transfected with V5-PIAS1 and E2. WB analysis demonstrates that E2 is co-IPed with V5-PIAS1. Input: whole-cell lysates probed for V5-PIAS1, E2, and β-actin (loading control). (C) Precipitated 3x-FLAG LANA by anti-FLAG magnetic bead were subjected to *in vitro* SUMOylation reaction using E1, E2, SUMO2, and PIAS1. WB analysis was performed with anti-Flag-HRP antibody. Bracket denotes SUMOylated LANA. (D) Purified V5-E2 were subjected to SUMOylation using E1, E2, SUMO2, PIAS1, and HPV-16 *oriP*-containing plasmid, pOri16M. WB analysis was performed with anti-V5-HRP antibody. Bracket denotes SUMOylated E2.

In summary, we identify PIAS1 as a key E3 SUMO ligase for EBNA1, enriched at EBV *oriP* where it binds and colocalizes with EBNA1. We uncover two novel SUMOylation sites (K17 and K75) along with K241, which plays important roles in regulating EBV lytic replication and episome retention (**Fig. 11**). These findings reveal a novel role for PIAS1 in controlling EBV latency through EBNA1 SUMOylation and open new avenues to investigate whether similar mechanisms regulate KSHV LANA and HPV E2.

**Figure 11.**
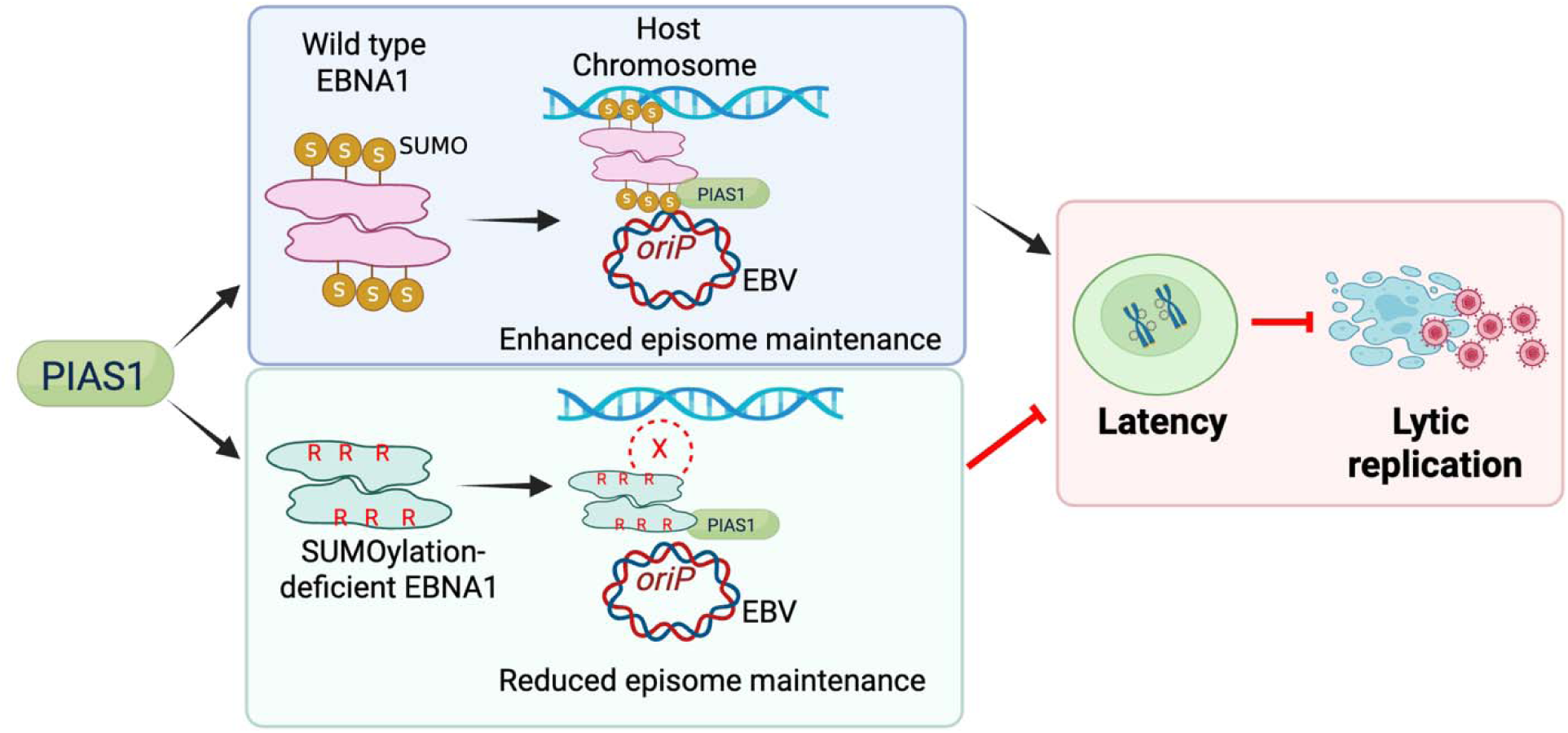
Model illustrating PIAS1-mediated SUMOylation of EBNA1 in EBV latency and genome maintenance. Wild-type EBNA1 undergoes PIAS1-mediated SUMOylation, which enhances tethering of the EBV genome to human chromosomes and stabilizes latent infection. Conversely, SUMOylation-deficient EBNA1 shows reduced genome tethering, thereby disrupting latency and facilitating lytic replication. Figure created with BioRender.com.

## MATERIAL AND METHODS

### Cell lines and cultures

Akata (EBV+) cells were cultured in Roswell Park Memorial Institute medium (RPMI 1640) supplemented with 10% FBS (Cat. # 26140079, Thermo Fisher Scientific) in 5% CO_2_ at 37°C [20, 43–46]. HEK-293 (EBV+) cells with B95.8 EBV genome was maintained in 150 μg/mL Hygromycin B (Cat. # J60681MC, Thermo Fisher Scientific). HEK-293 (EBV+), and 293T cells were cultured in Dulbecco’s modified eagle medium (DMEM) supplemented with 10% FBS in 5% CO_2_ at 37°C [47, 48]. See also **Table 1** for cell line sources.

**Table 1.**
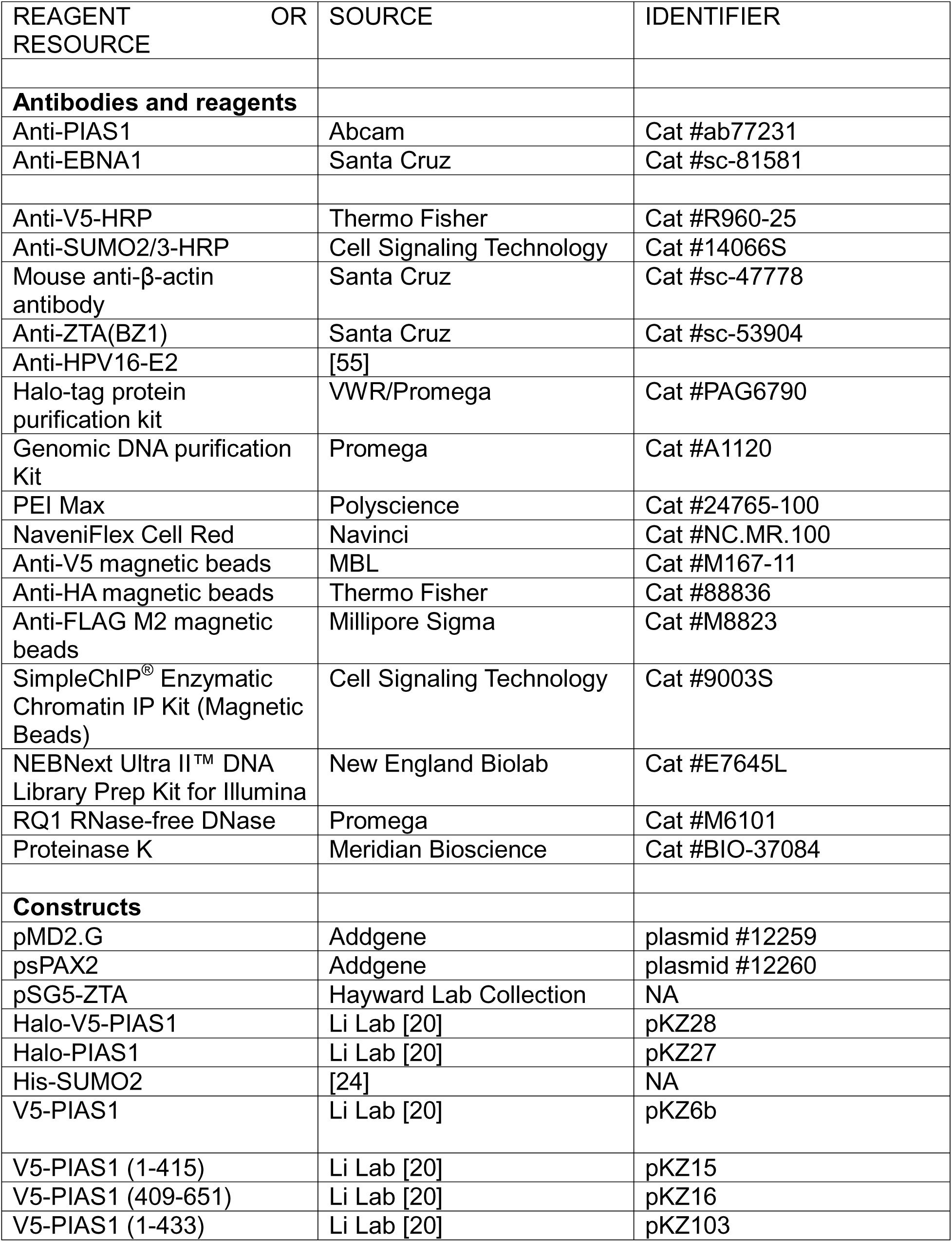
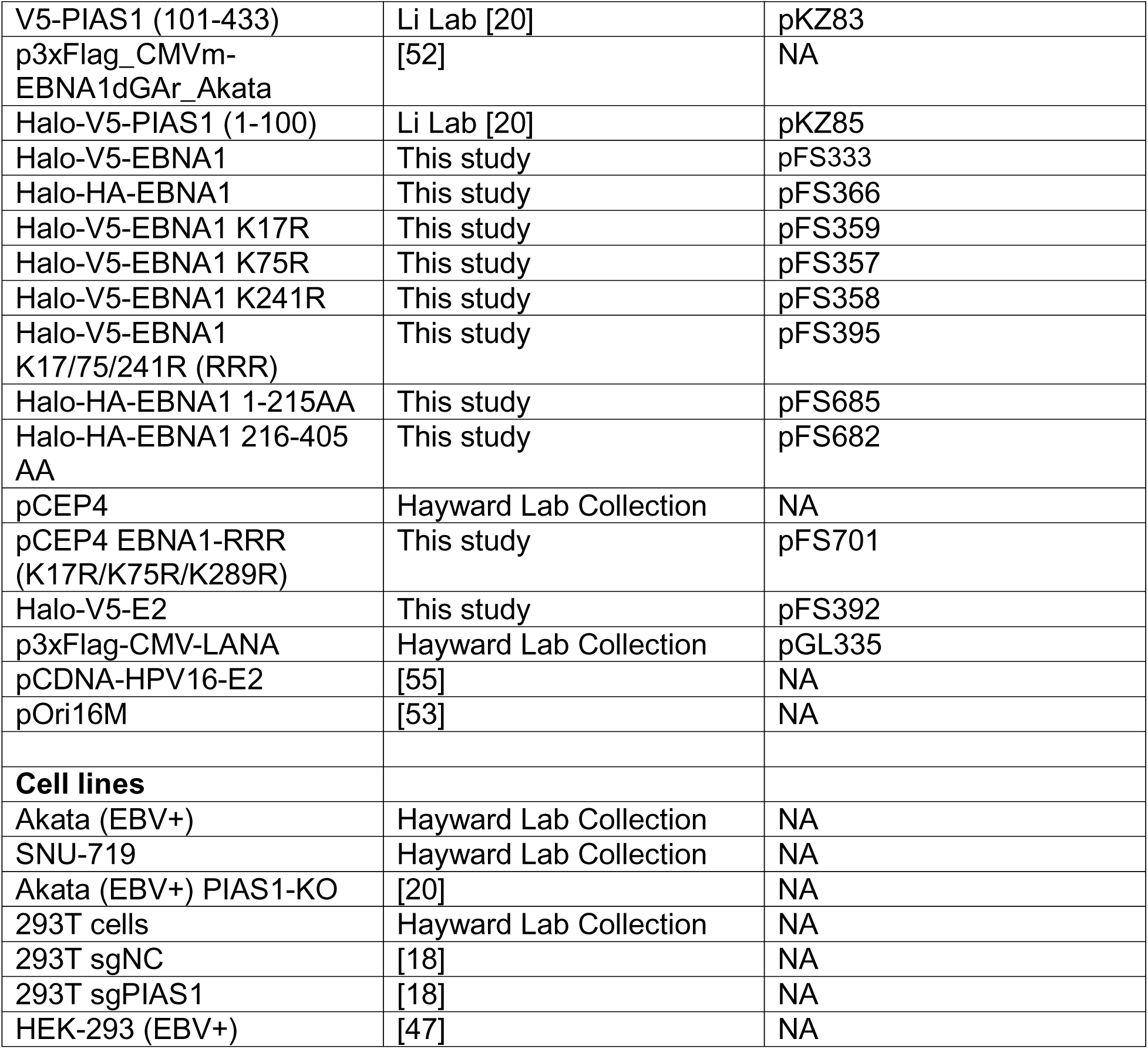
Antibody, reagents, constructs and cell lines.

### Chromatin-immunoprecipitation (ChIP) sequencing

Chromatin immunoprecipitation (ChIP) assays were performed using the SimpleChIP® Enzymatic Chromatin IP Kit (Magnetic Bead) (Cat. #9003S, Cell Signaling Technology) according to the manufacturer’s protocol. Briefly, 4–5 × 10 Akata (EBV+) cells were cross-linked with 1% formaldehyde and subsequently digested with micrococcal nuclease to yield chromatin fragments ranging from 150 to 900 bp. One percent of the digested chromatin was reserved as input control. For immunoprecipitation, 6 µg of chromatin was incubated with 1 µg of anti-PIAS1 antibody (Cat. #ab77231, Abcam). Following immunoprecipitation, cross-links were reversed, and DNA was purified for downstream analysis.

For ChIP-seq, 0.1–20 ng of ChIP-derived DNA was further sheared to an average fragment size of ∼200 bp using a Covaris system (2-minute sonication in 15 µL microTUBE-15 AFA Beads Screw-Cap tubes, Cat. #520145, Covaris). Sequencing libraries were prepared using the NEBNext® Ultra™ II DNA Library Prep Kit for Illumina (Cat. #E7645L, New England Biolabs), which features a low-input optimized adapter ligation system to reduce bias. Final libraries were quantified using both Qubit fluorometric quantitation and Bioanalyzer analysis, pooled, and subjected to next-generation sequencing on either the Illumina NovaSeq 6000 or NextSeq 2000 platforms.

### ChIP-Seq analysis

ChIP-Seq FASTQ files were processed and analyzed using the GALAXY web server [49]. Initially, raw reads were assessed for quality using FastQC, and low-quality sequences were trimmed accordingly. High-quality reads were then aligned to the human reference genome and the EBV Akata strain (GenBank: KC207813) using Bowtie2. Peak calling was performed with MACS2, using input DNA as a control to identify significant enrichment regions [50]. For data visualization, aligned reads and peaks were loaded into the Integrative Genomics Viewer (IGV) [51]. The resulting ChIP-Seq datasets have been deposited in the NCBI Gene Expression Omnibus (GEO) under accession number GSE306456 (**Reviewer token**: **ofujmaecvdmjzij**).

### ChIP-qPCR

ChIP process and qPCR were previously described [19]. DNA-protein complex were immunoprecipitated with anti-PIAS1 antibody (Cat. #ab77231, Abcam) and rabbit IgG control (Cat. #2729, Cell Signaling Technology). ChIP was performed using an Enzymatic Chromatin IP kit (Cell Signaling Technology, SimpleChIP Enzymatic Chromatin IP kit) as described previously [20]. IP-ed DNA was quantified by qPCR using *oriP*-specific primers (**Table S1**).

### Plasmid construction

Halo-PIAS1, Halo-V5-PIAS1 (full length and aa 1–100), and V5-PIAS1 (full length, aa 1–415, aa 409–651, and aa 101–433) plasmids were previously described [20].

EBNA1 dGAr insert was amplified from p3xFlag_CMVm-EBNA1 dGAr_Akata, a gift from Kathy Shair [52] by PCR with Q5 high-fidelity DNA polymerase (Q5-PCR) and cloned into the pHTN-CMV-Neo vector with an N-terminal V5 tag via Gibson Assembly. Halo-V5-EBNA1 was used as a template to create K17R, K75R, and K241R mutants using site-directed mutagenesis kit with Pfu Ultra II Fusion HotStart DNA Polymerase (Cat. #600672, Agilent) according to the manufacturer’s instructions on Quikchange II system (Cat. #200523, Agilent). The SUMOylation-deficient EBNA1 in pCEP4 was similarly mutated using the same approach. Halo-HA-EBNA1 (aa 1-215) was generated by adding stop codon after aa 215. For Halo-HA-EBNA1 (aa 216-405), the corresponding DNA was amplified by Q5-PCR and then digested using *EcoRI* and *NotI-HF* and cloned into pHTN-CMV-Neo vector.

HPV16-E2 insert was amplified from pCDNA-HPV16-E2 (a gift from Iain Morgan) by Q5-PCR and cloned into pHTN-CMV-Neo vector with an N-terminal V5-tag via Gibson Assembly. See also **Table 1** for construct sources. All primer sequences are listed in **Table S1**.

### *In situ* Proximity Ligation Assay (PLA)

PLA was modified as previously described [18], Briefly, cells were blocked with 3% BSA in PBS at room temperature for 1 h, then incubated with PBS control or a mixture of mouse anti-EBNA1 (Cat. #sc-81581, Santa Cruz) and rabbit anti-PIAS1 (Cat. #ab77231, Abcam) antibodies (1:50 dilution in 3% BSA) at 4°C overnight. Then the probes were incubated at 37°C for 1 h, followed by ligation and amplification (NaveniFlex Cell Red #NC.MR.100, Navinci). Cell nuclei were stained using Duolink *in situ* mounting media with 4′,6-diamidino-2-phenylindole (DAPI) and visualized by Nikon AXR confocal microscope.

### Plasmid retention assay

HEK-293T (non-targeting control and sg-PIAS1) cells were established previously [18]). 2μg pCEP4 (WT EBNA1 or SUMOylation-deficient mutant) was transfected into the cells using PEI max for 24h and the culture media was changed and the cells were selected under 150 μg/mL hygromycin B until stable cell lines were established around 14 days post transfection [34].

The cells were split into 3x10^5^ cells/mL in 10 cm plate with 10 mL of DMEM+10% FBS and incubated in 5% CO_2_ at 37°C without hygromycin B. One portion of the cells were harvested for WB analysis. Every 3 days, the cells were passaged and reseeded at the same density (3x10^5^ cells/mL) in fresh 10 cm plates with 10 mL of DMEM+10% FBS. This process was repeated for a total of 9 days. At each passage, 6x10^5^ cells were harvested for pCEP4 DNA detection. Total genomic DNA was extracted using the Genomic DNA Purification Kit (Cat. #A1120, Promega). Relative plasmids copy numbers were similarly measured by qPCR using previously described primers [34] and normalized with β-actin gene (**Table S1**).

### Cell lysis, immunoblotting and immunoprecipitation

Cell lysis, immunoprecipitation and immunoblotting (WB) were performed as previously described [18], with minor modifications. Cells were harvested, lysed in 2× SDS-PAGE sample buffer, and boiled for 5 min. Proteins were resolved on 4–20% TGX gels (Cat. #4561096; Bio-Rad), transferred to PVDF membranes, and probed with the indicated primary antibodies followed by horseradish peroxidase–conjugated secondary antibodies. See also **Table 1** for antibody sources.

For immunoprecipitation, cells were lysed in buffer containing 50 mM Tris-HCl (pH 7.5), 150 mM NaCl, 0.1% NP-40, and protease inhibitor cocktail (Cat. #4693116001; Sigma-Aldrich) on ice for 30 min. Lysates were sonicated (10 seconds on/10 seconds off, three cycles, 35% amplitude) and clarified by centrifugation at 14,600 × g for 15 min at 4 °C. Ten percent of the supernatant was reserved as input, and the remainder was incubated with the indicated magnetic beads. Input and immunoprecipitated proteins were analyzed by immunoblotting by the indicated antibodies. Anti-HPV16-E2 antibody is a gift from Iain Morgan.

### Protein expression and purification

Halo-tagged PIAS1, EBNA1, and HPV-16 E2 proteins were expressed and purified as previously described [18], with minor modifications. Briefly, HEK293T cells were transfected with 18 μg plasmid DNA and 54 μg PEI Max and harvested 48 h later. Cells were lysed in Halo purification buffer (50 mM HEPES, pH 7.5, 150 mM NaCl, 1 mM EDTA, 0.005% NP-40, 1 mM DTT), sonicated (10 seconds on/10 seconds off, three cycles, 35% amplitude), and clarified by centrifugation at 14,600 × g for 15 min at 4 °C. The supernatant was incubated with 200 μL pre-washed Halo resin at 4 °C overnight. Beads were washed three times with Halo purification buffer and subsequently incubated with Halo purification buffer containing TEV protease at 4 °C overnight.

### *In vitro* SUMOylation assay

*In vitro* SUMOylation assay was performed using the SUMO2 conjugation kit as previously described [18, 19], with minor modifications. Reactions were carried out in buffer containing 40 mM Tris (pH 7.1), 40 mM NaCl, 1 mM β-mercaptoethanol, and 5 mM MgCl₂. The substrates (EBNA1, LANA or HPV-16 E2) were incubated with 100 nM SAE1/SAE2 (E1), 2 μM His₆-Ube2I/UBC9 (E2), 50 μM SUMO2, and 4 mM ATP, with PIAS1 as the E3 ligase in the presence or absence of 50 ng/μL pCEP4 or pOri16M [53] plasmid. Reactions were incubated at 37 °C for 3 h, and SUMOylation was assessed by immunoblotting.

For *in vitro* SUMOylation of LANA, HEK293T cells were transfected with 10 μg of p3XFLAG-CMV-LANA, plasmid from Diane Hayward’s Lab collection, and harvested 48 h post-transfection. Cells were lysed in lysis buffer, and LANA was immunoprecipitated using Anti-FLAG M2 magnetic beads (Cat. #M8823; Millipore Sigma). SUMOylation reactions were performed directly on the bead-bound protein with gentle agitation.

### Lytic induction and EBV copy number detection

For lytic induction of EBV in HEK-293 (EBV+) cells, the cells were transfected with EBV ZTA plus other plasmids as indicated using PEI max for 48 h as described previously [18]. Extracellular viral DNA was extracted and quantified following established protocols [18, 22]. Briefly, EBV-containing media was treated with RQ1 RNase-free DNase (Cat. #M6101; Promega) to remove naked DNA, and the reaction was terminated with the supplied stop buffer. Proteinase K (Cat. #BIO-37084; Meridian Bioscience) and SDS were then added to digest viral proteins and to release virion-associated DNA. EBV DNA was purified by phenol–chloroform extraction and precipitated with isopropanol, sodium acetate, and glycogen at −80 °C overnight. DNA pellets were washed with 70% ethanol, air-dried, and resuspended in Tris-EDTA (TE) buffer (10mM Tris and 1mM EDTA, pH 8.0). EBV DNA was detected by PCR using BALF5-specific primers [18].

### Structure prediction by AlphaFold3

AlphaFold3 algorithm [25] was employed to predict the three-dimensional structure of EBNA1 (dGAr) dimer with 1XFR DNA sequence (5’- GGATAGCATATACTACCCGGATATAGATTA-3’). Molecular graphics of EBNA1 were performed with UCSF ChimeraX [54], developed by the Resource for Biocomputing, Visualization, and Informatics at the University of California, San Francisco, with support from National Institutes of Health R01-GM129325 and the Office of Cyber Infrastructure and Computational Biology, National Institute of Allergy and Infectious Diseases. Model 1 of each prediction was used to display EBNA1 structure and PIAS1 interaction with EBNA1.

### Electrophoretic Mobility Shift Assay

Purified V5-EBNA1 WT and EBNA1 RRR proteins were serially diluted (0–500 nM) and incubated with a 2X FR DNA probe labeled with IRDye 700 (10 nM) [5] for 3h at 30°C. Reactions were stopped by the addition of 2X nucleic acid loading buffer (50 mM Tris-HCl pH 8.0, 20% glycerol, 2 mM EDTA, 0.1% Bromphenol Blue) and resolved on a 1.4% agarose gel at 80 V in 1X TBE running buffer. DNA–protein complexes were visualized using a LI-COR Odyssey Fc imaging system under the 700 nm channel.

### Quantification and statistical analysis

Statistical analyses were performed using a two-tailed Student *t*-test with Microsoft Excel software. A *p* value less than 0.05 was considered statistically significant. The values are presented as means and standard deviations for biological replicate experiments as specified in the figure legends. The Fig.11 was created using BioRender.

## ACKNOWLEDGEMENTS

We thank S. Diane Hayward (Johns Hopkins University) for providing reagents, plasmids, and cell lines. We are grateful to Didier Trono (EPFL) for the pMD2.G and psPAX2 plasmids (Addgene plasmid nos. 12259 and 12260); Ronald Hay (University of Dundee) for the His-SUMO2 plasmid; Kathy Shair (University of Pittsburgh) for the p3xFLAG_CMVm-EBNA1dGAr_Akata plasmid; Iain Morgan for the pOri16M, HPV16 E2 plasmid and E2 antibody. We also thank Henri-Jacques Delecluse (German Cancer Research Centre) and Ayman El-Guindy (Yale University) for providing HEK293 cells carrying B95.8 EBV genomes. We also thank Haitao Guo (University of Pittsburgh) for generously providing access to the Li-Cor imaging system used for EMSA gel visualization.

This work was supported in part by grants from the National Institute of Allergy and Infectious Diseases (AI141410 and AI187186) awarded to R.L. R.L. also received a Research Scholar Grant (134703-RSG-20–054-01-MPC) from the American Cancer Society, as well as support from the University of Pittsburgh Medical Center, Hillman Cancer Center, Virginia Commonwealth University (VCU) Philips Institute for Oral Health Research, and the VCU Presidential Quest for Distinction Award. The funders had no role in study design, data collection and analysis, decision to publish, or preparation of the manuscript.

Conceptualization: R.L., F.G.S.;

data curation: F.G.S., K.Z., and R.L.;

formal analysis: F.G.S. and R.L.;

funding acquisition: R.L.;

investigation: F.G.S., K.Z., and R.L.;

methodology: F.G.S., K.Z., and R.L.

project administration: R.L.;

resources: R.L.;

supervision: R.L.;

validation: F.G.S., K.Z., and R.L.;

visualization: F.G.S., K.Z., and R.L.;

writing (original draft): F.G.S.;

writing (review and editing): R.L. and F.G.S.

